# A screen for synthetic genetic interactions with the *Saccharomyces cerevisiae hrq1ΔN* allele

**DOI:** 10.1101/2025.07.03.663020

**Authors:** Kennadi A. Shumaker, Michael E. Kumcu, Faith E. McDevitt, Cody M. Rogers, Matthew L. Bochman

## Abstract

The *Saccharomyces cerevisiae* Hrq1 helicase is a functional homolog of the disease-linked human RECQL4 enzyme and has been used as a model to study RecQ4 helicase subfamily biology. Although the motor cores of Hrq1 and RECQL4 are quite similar, these proteins display distinct N-terminal domains (NTDs) of unknown function. Do these domains facilitate species-specific activities by the two helicases, or do they serve common roles despite their differences in sequence and predicted structure? We probed these questions here by analyzing an NTD-truncated isoform of Hrq1 (Hrq1ΔN) both *in vitro* and *in vivo*. We found that the Hrq1 NTD houses a cryptic DNA binding site and that the *hrq1ΔN* allele is distinct from both *hrq1Δ* and the catalytically inactive *hrq1-K318A* mutant. Using synthetic genetic array analysis of *hrq1ΔN* crossed to the yeast *S. cerevisiae* single-gene deletion and temperature-sensitive allele collections, we also identified hundreds of synthetic genetic interactions. As with similar analyses of *hrq1Δ* and *hrq1-K318A*, our results suggest roles for Hrq1 and its NTD in multiple physiological pathways that underpin genome integrity. Together, these data are guiding our ongoing efforts to understand the roles of Hrq1 and RECQL4 in genome maintenance, which will help to explain why RECQL4 mutations cause disease.

**ARTICLE SUMMARY:** This work should interest researchers in the genome integrity and yeast disease model fields. It continues ongoing efforts to develop the *Saccharomyces cerevisiae* Hrq1 helicase as a model to understand the disease-linked human RECQL4 helicase. Hrq1 and RECQL4 share similar helicase cores but have divergent N-terminal domains (NTDs) of unknown function. The results demonstrate that the Hrq1 NTD contains a DNA binding site, and an *HRQ1* allele lacking its NTD (*hrq1ΔN*) interacts with hundreds of mutants in defined allele collections. Thus, the *hrq1ΔN* allele is distinct from the previously characterized *hrq1Δ* and *hrq1-K318A* (catalytically inactive) mutants.

## INTRODUCTION

Mutations in the human RECQL4 DNA helicase cause three autosomal recessive diseases of genomic instability: Rothmund-Thomson (Ho, Tong et al. 2025), Baller-Gerold (Van Maldergem, Piard et al. 1993), and RAPADILINO syndromes (Vargas, de Almeida et al. 1992). Although all distinct, these diseases do include some overlapping pathologies, including bone morphology defects and predispositions to cancers (Balajee 2021, Xu, Chang et al. 2021). However, it is unclear how mutations in the helicase ultimately precipitate a disease state. Part of this quandary is the fact that RECQL4 is an evolutionary chimera, appearing to be a fusion of an essential DNA replication initiation factor and a RecQ family helicase (Liu 2010). The majority of the disease-linked alleles are located in the helicase portion of the protein (Larizza, Roversi et al. 2010), presumably because perturbing the replication initiation function of RECQL4 would be embryonic lethal, but it is difficult to study the effects of these mutations without pleiotropically impacting DNA replication. Biochemical investigations of RECQL4 are similarly stymied by the protein’s large size (∼150 kDa) and natively disordered N-terminal domain (NTD), making recombinant protein purification difficult (Bochman, Paeschke et al. 2014, Keller, Kiosze et al. 2014, Rogers, Wang et al. 2017), as well as its strong annealing activity masking its DNA unwinding activity (Macris, Krejci et al. 2006, Xu and Liu 2009, Keller, Kiosze et al. 2014).

To overcome these technical hurdles, we and others study the *Saccharomyces cerevisiae* Hrq1 helicase, which is homologous to the RecQ portion of RECQL4 but lacks the confounding NTD related to replication initiation (Barea, Tessaro et al. 2008, Bochman, Paeschke et al. 2014, Choi, Min et al. 2014, Rogers and Bochman 2017, Rogers, Wang et al. 2017, Nickens, Rogers et al. 2018, Nickens, Sausen et al. 2019, Rogers, Lee et al. 2020, Rogers, Sanders et al. 2020, Sanders, Nguyen et al. 2020, Nickens and Bochman 2021, Luong, Li et al. 2022, Nickens and Bochman 2022, Rajapaksha, Simmons et al. 2022, Nickens, Feng et al. 2024). To date, this work has drawn functional parallels between Hrq1 and RECQL4, highlighting roles in genome maintenance processes such as DNA repair (Bochman, Paeschke et al. 2014, Rogers, Wang et al. 2017, Rogers, Lee et al. 2020, Rogers, Simmons III et al. 2020, Luong, Li et al. 2022) and telomerase regulation (Bochman, Paeschke et al. 2014, Nickens, Rogers et al. 2018, Nickens, Sausen et al. 2019, Nickens and Bochman 2021, Nickens, Feng et al. 2024). However, like human RECQL4, *S. cerevisiae* Hrq1 contains a large NTD of unknown function.

Are there cryptic nucleic acid binding sites in the Hrq1 NTD as observed for RECQL4 (Keller, Kiosze et al. 2014)? Are there replication-independent roles shared by the Hrq1 and RECQL4 NTDs? Are there species-specific functions? We sought to answer these questions here by characterizing an N-terminal truncation of residues 1-279 of Hrq1 (Hrq1ΔN) both *in vitro* and *in vivo*. As part of the latter work, we performed synthetic genetic array (SGA) analysis of the *S. cerevisiae hrq1ΔN* allele by crossing it to the single-gene deletion collection (Giaever and Nislow 2014) and the temperature-sensitive (TS) collection (Kofoed, Milbury et al. 2015) to generate a comprehensive set of double-mutant strains for analysis. This will allow us to compare synthetic genetic interactions with *hrq1ΔN* to those observed for *hrq1Δ* and *hrq1-K318A* (Rogers, Sanders et al. 2020, Sanders, Nguyen et al. 2020) to determine NTD-specific effects.

## METHODS & MATERIALS

### Media and other reagents

*S. cerevisiae* and *Escherichia coli* cells were grown in standard media at 30 and 37°C, respectively, unless otherwise noted. Radioisotopes were purchased from PerkinElmer, and unlabeled ATP was from GE Healthcare. All restriction enzymes were from NEB, and all oligonucleotides were from IDT. All other reagents used were of the highest grade available.

### Recombinant protein production

Full-length Hrq1 was over-expressed in baculovirus-infected insect cell culture and purified by Ni-affinity chromatography as previously described (Rogers, Wang et al. 2017). The Hrq1ΔN expression plasmid was created by PCR-amplifying the coding sequence for Hrq1 amino acids 280-1077 from pMB309 (Bochman, Paeschke et al. 2014) with oligonucleotides MB635 (CATGCATGTCATGAGCTGGTCTCATCCACAATTCGAAAAGGGTGCCAACTGGTCTCATCCACAATTCGAAAAGGGTGCCAACTGGTCTCATCCACAATTCGAAAAGGGTGGTGGAGGCGGTGGTGGAGGTTTGGAAGTTTTGTTCCAAGGTCCAGGATCCCCGGAAGTATATCAGGGTATGGAACACG) and MB530 (CGTACGCTCGAGTATCTCTTTTTTAATGATTTCATTTGTATGAGTATCGTCTTTCGTAGC), digesting the PCR product with *Bsp*HI and *Xho*I, and ligating it into the *Nco*I and *Xho*I sites of pET21d (Novagen) to create pMB296. The Hrq1 N-terminal domain (NTD) expression plasmid was generated by PCR-amplifying the coding sequence for Hrq1 amino acids 1-279 from pMB309 using oligonucleotides MB653 and MB1032 (CGATCGATCTCGAGTGCAAGCTCAAAACAGAGACCTTTATACTTCGC), digesting the PCR product with *Bsp*HI and *Xho*I, and ligating it into the *Nco*I and *Xho*I sites of pMB131 (Paeschke, Bochman et al. 2013, Andis, Sausen et al. 2018) to create pMB426. These plasmids were transformed into Rosetta^TM^ 2(DE3) pLysS cells (Novagen), and the Hrq1 truncations were over-expressed using the high-density autoinduction method (Studier 2005). Both proteins were purified using Strep-Tactin Sepharose and Ni-affinity chromatography as previously described for full-length Hrq1 (Bochman, Paeschke et al. 2014). Additional details concerning the cloning of the expression constructs and protein purification schemes are available upon request.

### Transmission electron microscopy (TEM)

To prepare negatively stained grids, 4 μL of protein solution (∼20 ng/μL) was applied to glow-discharged 300-mesh carbon-coated copper grids (Electron Microscopy Sciences). Excess solution was removed by blotting with filter paper, and subsequently, the grid was stained with 0.75% uranyl formate (Electron Microscopy Sciences) and allowed to air dry. The grids were then imaged under low-dose conditions (≤20 e^-^/Å^2^) using a JEOL 300-kV 3200FS transmission electron microscope with the energy filter operated at an energy slit of 20 eV to minimize potential beam-induced damage to the protein samples. The digital images were collected using a Gatan 4k x 4k CCD camera at a nominal magnification of 80,000x, yielding a 1.5 Å/pixel resolution in the final images.

### *In silico* protein analyses

AlphaFold 3 models were rendered on the Google DeepMind AlphaFold Server (<u>alphafoldserver.com</u>) using default parameters, unless otherwise noted. The protein sequence for *S. cerevisiae* Hrq1 from the S288c genetic background was obtained from the *Saccharomyces* Genome Database (yeastgenome.org). The *Schizosaccharomyces pombe* Hrq1 sequence was from PomBase (https://www.pombase.org/gene/SPAC23A1.19c). The other RecQ4 family helicase sequences were retrieved from UniProt (https://www.uniprot.org/). See Table S1 for additional details. Structures were visualized and analyzed in Chimera X (https://www.cgl.ucsf.edu/chimerax). The disorder analyzes utilized online PrDOS (https://prdos.hgc.jp/cgi-bin/top.cgi), DEPICTER (https://biomine.cs.vcu.edu/servers/DEPICTER/), DISOPRED3 (https://github.com/psipred/disopred), and IUPred (https://iupred2a.elte.hu/) servers with default parameters.

### Diepoxybutane (DEB) sensitivity assay

Wild-type MBY49 (*MATa ura3-52 lys2-801_amber ade2-101_ochre trp1Δ63 his3Δ200 leu2Δ1 hxt13::URA3*) (Paeschke, Bochman et al. 2013, Bochman, Paeschke et al. 2014) and its *hrq1Δ* (MBY161, *hrq1::his3MX6*), *hrq1-K318A* (MBY346, *hrq1::hrq1-K318A(natMX)*), and *hrq1ΔN* (MBY462, *hrq1::hrq1ΔN(natMX)*) derivatives were tested for DEB sensitivity essentially as described (Bochman, Paeschke et al. 2014). Briefly, the cells were grown overnight in 1% yeast extract, 2% peptone, and 2% dextrose (YPD) medium at 30°C with aeration, diluted to an optical density at 600 nm of 1 with sterile water, and then 10-fold serially diluted likewise to 10^-4^.

Then, 5 µL of each dilution was spotted onto plates either containing or lacking DEB (Sigma) at the indicated concentration. The plates were incubated in the dark at 30°C for ∼2 d before being imaged with a flat-bed scanner. DEB sensitivity assays were performed in triplicate with representative scans presented in Figure 3. All strains were derived from the wild-type YPH genetic background (Sikorski and Hieter 1989) by standard methods. Details of the strain constructions are available upon request.

### NADH-coupled ATPase assay

ATPase reactions were performed in ATPase buffer (25 mM Na-HEPES [pH 8.0], 5% glycerol, 50 mM NaOAc [pH 7.5], 150 µM NaCl, and 0.01% NP-40 substitute) including the following reagents: 5 mM ATP (pH 7.0) (DOT Scientific), 5 mM MgCl_2_, 0.5 mM phosphor(enol)pyruvic acid (Sigma), 0.4 mM NADH (MP Biomedicals, LLC), 5 U/mL rabbit pyruvate kinase (Roche), and 8 U/mL lactate dehydrogenase from oyster (Sigma). Unless otherwise stated, the helicase concentration was 10 nM in all reactions, and poly(dT)_50_ single-stranded (ss)DNA was used at 1 μM. To minimize possible confounding effects caused by secondary structures that the oligonucleotides may form, the substrates were diluted to their working concentrations, boiled, and snap-cooled prior to addition to the ATPase assays. Absorbance at 340 nm was read at 37°C in 96-well plates using a BioTek Synergy H1 microplate reader. Absorbance readings were converted to ATP turnover based on NADH concentration. It was assumed that 1 µM NADH oxidized is proportional to 1 µM ATP hydrolyzed.

### DNA binding

DNA binding was measured using electrophoretic mobility shift assays (EMSAs). The proteins were incubated at the indicated concentrations with 0.1 nM radiolabeled DNA for 30 min at 30°C in binding buffer (25 mM Na-HEPES [pH 8.0], 5% glycerol, 50 mM NaOAc [pH 7.6], 150 µM NaCl, 7.5 mM MgOAc, and 0.01% Tween-20). Protein-DNA complexes were separated from unbound DNA on 8% 19:1 acrylamide:bis-acrylamide gels in 1x TBE (90 mM Tris–HCl [pH 8.0], 90 mM boric acid, and 2 mM EDTA [pH 8.0]) buffer at 10 V/cm. Gels were dried under vacuum and imaged using a Typhoon 9210 Variable Mode Imager. DNA binding was quantified using ImageQuant 5.2 software. The DNA substrate was made by 5ʹ-end labelling a poly(dT) 30mer oligonucleotide with T4 polynucleotide kinase (T4 PNK; NEB) and γ[^32^P]-ATP. Labelled oligonucleotides were separated from free label using illustra ProbeQuant G-50 micro columns (GE Healthcare) following the manufacturer’s instructions.

### SGA analysis

To generate the *hrq1ΔN* SGA query strain, the natMX6-tagged allele was PCR-amplified from MBY462 using oligonucleotides MB527 (GTGAATTGCTCAGAAGAGAAAGGCATACCGTC) and MB528 (CTGTGCATCAACAAGGTGACAGAATGTTGATG). This PCR product was then used to transform strain Y8205 (*MATα can1Δ ::STE2pr-Sp_his5 lyp1Δ ::STE3pr-LEU2 his3Δ1 leu2Δ0 ura3Δ0*) (Tong, Evangelista et al. 2001, Tong, Lesage et al. 2004). The replacement of the endogenous *HRQ1* gene with the *hrq1ΔN* allele was verified by PCR analysis using genomic DNA and oligonucleotides MB527 and MB528 that anneal to regions up- and downstream of the *HRQ1* locus. The confirmed *hrq1ΔN* strain was named MBY641.

SGA analysis of the *hrq1ΔN* mutant was performed at the University of Toronto using previously described methods (Tong, Evangelista et al. 2001, Tong, Lesage et al. 2004). MBY641 was crossed to both the *S. cerevisiae* single-gene deletion collection (Giaever and Nislow 2014) and the TS alleles collection (Kofoed, Milbury et al. 2015) to generate double mutants for analysis. For these screens, the control strain (Y8835) contained the NatMX marker inserted into the benign *ura3* locus (*MATα ura3Δ::natMX4 can1Δ::STE2pr-Sp_his5 lyp1Δ his3Δ1 leu2Δ0 ura3Δ0 met15Δ0 LYS2+*). Quantitative scoring of the genetic interactions was based on colony size. The SGA score measures the extent to which a double-mutant colony size deviates from the colony size expected from combining two mutations together. The data include both negative (putative synthetic sick/lethal) and positive interactions (potential epistatic or suppression interactions) involving *hrq1ΔN*. The magnitude of the SGA score is indicative of the strength of the interaction. Based on statistical analysis, it was determined that a default cutoff for a significant genetic interaction is *p* < 0.05 and SGA score > |0.08|. It should be noted that only top-scoring interactions were confirmed by remaking and reanalyzing the double mutants by hand.

### Confirmation of top SGA hits

The top five positive and negative interactors with *hrq1ΔN* from the single-gene deletion and TS arrays were reanalyzed by hand to confirm their phenotypes. Briefly, the SGA query strain MBY641 (Nat^R^) was mated to *MATa* tester strains from the arrays (Kan^R^), sporulated, and then analyzed by random spore analysis (Lichten 2014), spot dilution (Andis, Sausen et al. 2018), and/or growth curve (Ononye, Sausen et al. 2020, Sausen and Bochman 2021) assays of Nat^R^ Kan^R^ spore clones compared to the parental single-mutant strains and wild-type.

### Statistical analysis

Data were analyzed and graphed using GraphPad Prism 6 software. The reported values are averages of ≥ 3 independent experiments, and the error bars are the standard deviation. *P*-values were calculated as described in the figure legends, and we defined statistical significance as *p* < 0.01.

## RESULTS AND DISCUSSION

### The Hrq1 NTD is predicted to contain native disorder

The N-terminus of human RECQL4 is intrinsically disordered yet contains important functional elements (Keller, Kiosze et al. 2014). Although the Hrq1 NTD does not display similarity to the RECQL4 NTD at the primary sequence level, we hypothesized that the native disorder might be conserved rather than the amino acid sequence. We base this conjecture on the fact that negatively stained TEM images of single Hrq1 particles have a multi-lobed ‘C-’ or ‘O-shaped’ structure (Fig. 1A and (Rogers, Wang et al. 2017)) that is roughly consistent with the size and shape of the AlphaFold-predicted structure of Hrq1 minus its N-terminal ∼279 amino acids (Fig. 1A). In addition to this portion of the structure being predicted to include long stretches of random coil, the confidence of the prediction is lower in the NTD compared to the helicase core of the enzyme, consistent with a propensity for molecular motion or disorder (Abramson, Adler et al. 2024). This predicted combination of intrinsic disorder and low-confidence secondary structural elements is also displayed by the AlphaFold model of the *Schizosaccharomyces pombe* Hrq1 (Fig. 1B), though in the *Drosophila melanogaster* RecQ4 (Fig. 1C) and human RECQL4 (Fig. 1D) homolog models, the amount of random coil is increased. Figure S1 shows additional AlphaFold models of RecQ4 family helicases and their NTDs.

**Figure 1.**
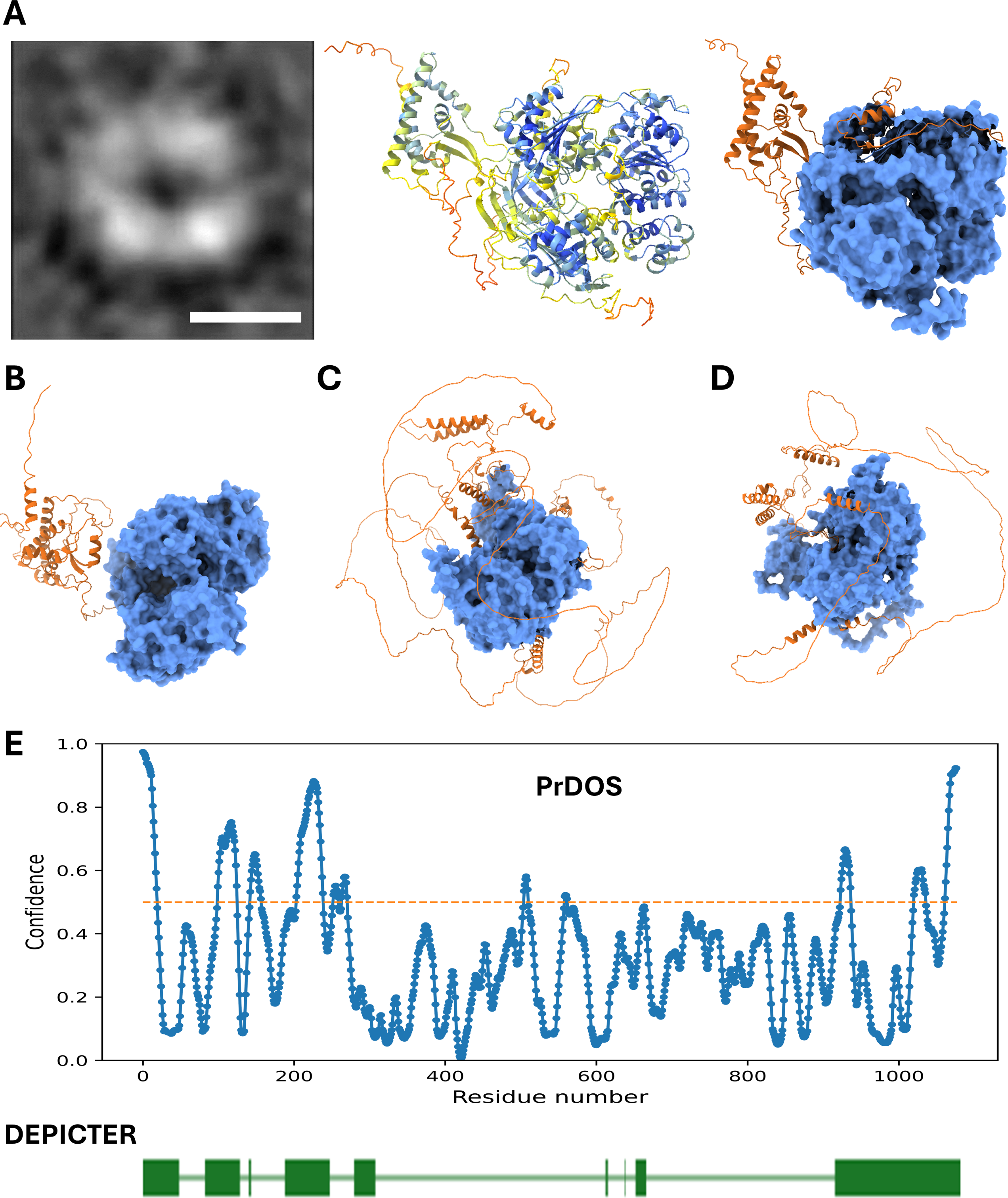
Structural considerations for the N-terminal domain (NTD) of Hrq1. **A)** Transmission electron micrograph of a recombinant Hrq1 particle (left; scale bar = 100 Å), an AlphaFold 3 prediction of the Hrq1 structure colored by confidence score (middle; cooler colors = more confident), and the same predicted structure depicted with the helicase core space filled and the NTP (aa 1-279) in ribbon format (right). These images depict a toroidal structure with a central void, but density corresponding to the NTD is missing in the micrograph, suggesting that the NTD is highly conformationally flexible or intrinsically disordered. AlphaFold predictions for *S. pombe* Hrq1 **(B)**, *D. melanogaster* RecQ4 **(C)**, and human RECQL4 **(D)** are also show. **E)** Disorder prediction for Hrq1 indicates a high confidence for native disorder in the NTD. The PrDOS server (top) predicts several disordered regions in the NTD (residues with values above the dotted line). The DEPICTER server (bottom) also predicts native disorder in the NTD (green boxes), as well as a largely disordered C-terminus.

To further probe the predicted structure of the Hrq1 NTD, we submitted the Hrq1 sequence to the PrDOS (Ishida and Kinoshita 2007) and DEPICTER (Barik, Katuwawala et al. 2020) protein disorder prediction servers for analysis. As shown in Figure 1E, both algorithms predict several regions of native disorder in the Hrq1 NTD (as well as the C-terminus), though the predictions do not match in the number or length of the disordered motifs. A similar analysis using the DISOPRED (Jones and Cozzetto 2015) and IUPred (Erdos and Dosztanyi 2024) servers was also performed but with similarly inconsistent results (Fig. S2). The DISOPRED prediction closely resembles that of PrDOS (Fig. 1B and S2A), but the IUPred algorithm predicts intrinsic disorder only at the very N-terminal portion of the Hrq1 NTD (Fig. S2B).

Based on the structure prediction (Fig. 1A) and three of four disorder predictors (Fig. 1E and S2A), it appears likely that the N-terminus of Hrq1 contains regions of native disorder but also folded secondary structures. The NTD of the second *S. cerevisiae* RecQ family helicase Sgs1 is somewhat similar, with long stretches of disordered amino acids interrupted by short motifs that dynamically sample folded and unfolded space, as indicated by NMR analysis (Kennedy, Daughdrill et al. 2013). Importantly, these portions of Sgs1 are necessary for its *in vivo* function. Although we only consider mutation of Hrq1 in the context of a full NTD truncation herein, future research should be devoted to disrupting the predicted secondary structures in the Hrq1 NTD to determine their physiological roles as a finer-grained approach.

### The NTD affects Hrq1’s ATPase and DNA binding activities

To investigate the biochemical roles of the Hrq1 NTD, we generated recombinant full-length wild-type and Hrq1ΔN proteins. We previously demonstrated that Hrq1 and RECQL4 are DNA-stimulated ATPases, whose activities are impacted by DNA sequence and structure (Rogers and Bochman 2017, Rogers, Wang et al. 2017). Here, we tested the Hrq1ΔN preparation for ATPase activity and found that it was identical to that of full-length Hrq1 (Fig. 2A), suggesting that the helicase core of Hrq1ΔN folds properly in the absence of the NTD. However, the ∼5-fold stimulation of wild-type Hrq1 ATPase activity by ssDNA is lacking in the Hrq1ΔN preparation. Addition of the purified NTD in *trans* to the Hrq1ΔN ATPase assays had no effect, nor did the NTD display ATP hydrolysis on its own (data not shown). Taken together, these data suggest that the Hrq1 N-terminus has an important role in coordinating ssDNA binding and ATPase activity.

**Figure 2.**
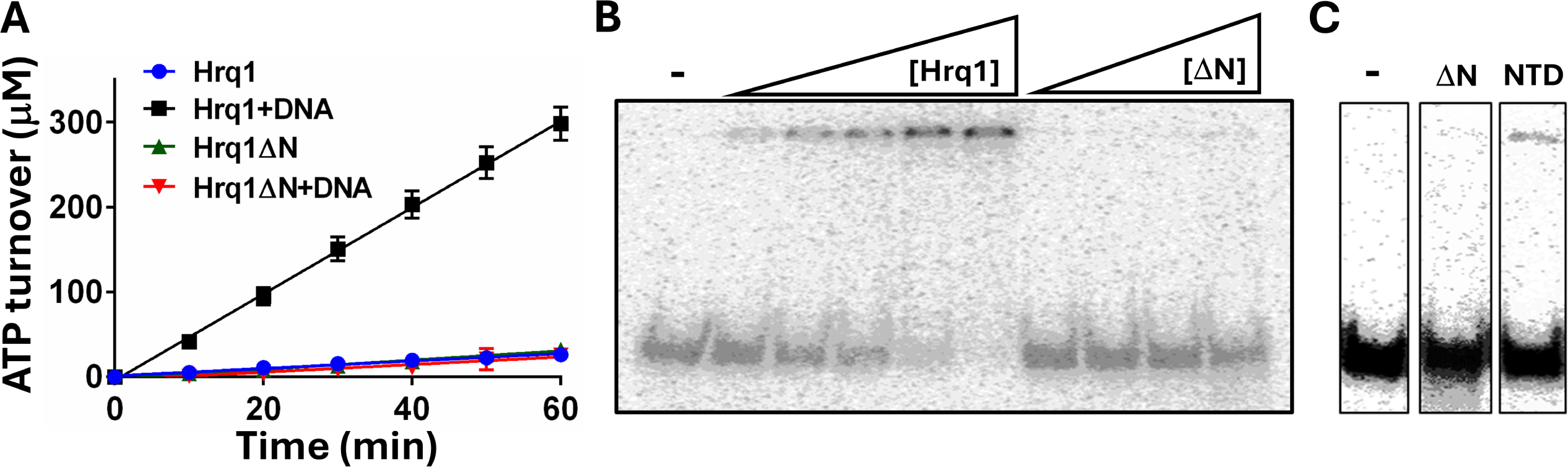
The Hrq1 NTD affects ATP hydrolysis and DNA binding. **A)** Hrq1 and Hrq1ΔN (10 nM) have low basal levels of ATPase activity. The addition of 1 μM poly(dT) ssDNA stimulates Hrq1 ATP hydrolysis (Hrq1+DNA) but has no effect on Hrq1ΔN (Hrq1ΔN+DNA). The plotted points are the averages of ≥3 independent experiments, and the error bars are the standard deviation. **B)** Gel shift analysis of Hrq1 (0.2-5 nM) and Hrq1ΔN (0.5-5 nM) binding to poly(dT) ssDNA. Hrq1ΔN is largely devoid of ssDNA binding activity, consistent with ssDNA failing to stimulate its ATPase activity. **C)** Example gel shift data showing that the Hrq1 NTD in isolation displays a weak ssDNA binding activity. -, no protein added; ΔN, 10 nM Hrq1ΔN added; and NTD, 10 nM Hrq1 NTD added.

**Figure 3.**
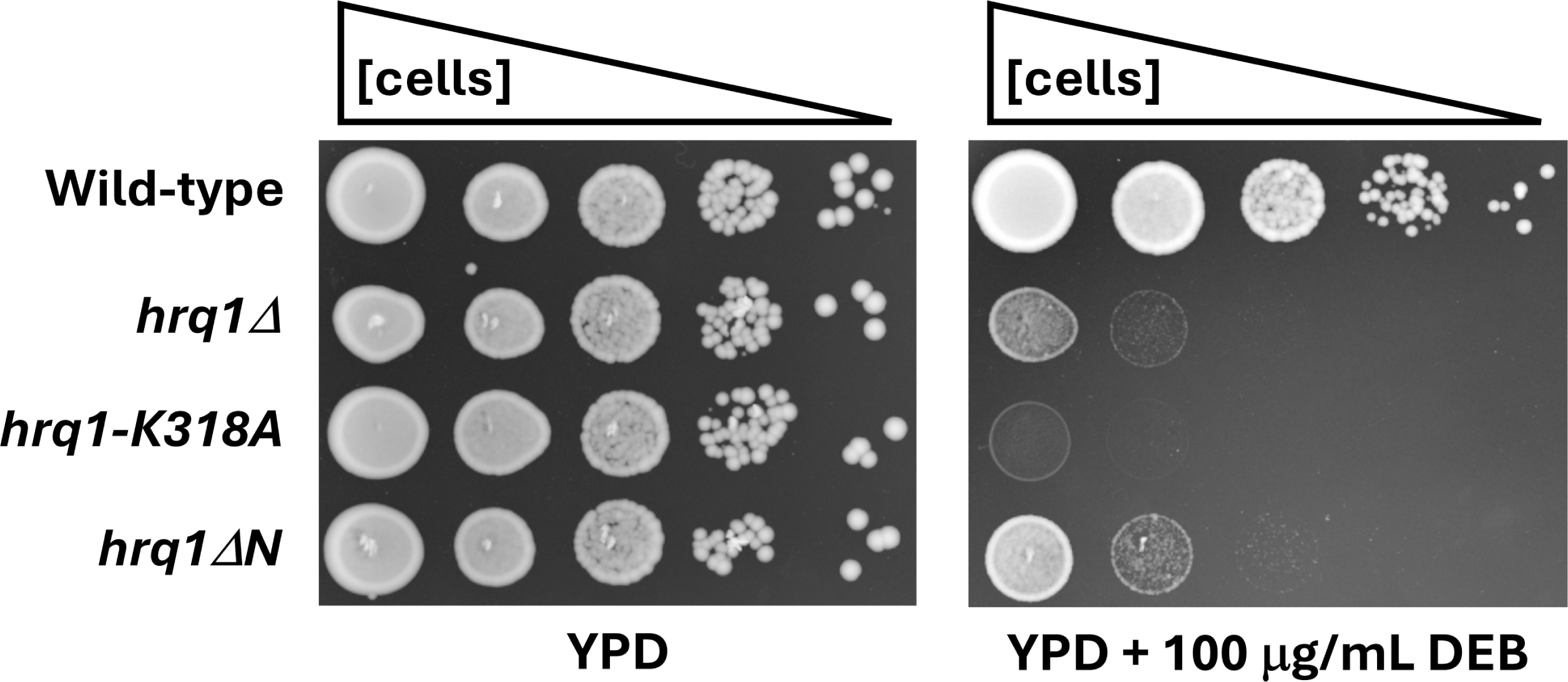
Cells expressing Hrq1ΔN phenocopy the diepoxybutane (DEB) sensitivity of *hrq1Δ* cells rather than the stronger sensitivity of *hrq1-K318A* cells. The plate images show the growth of 10-fold serial dilutions of the indicated strains at a uniform density incubated for 48 h at 30°C.

To test this directly, we performed ssDNA binding assays with Hrq1 and Hrq1ΔN. As previously demonstrated, gel shift assays showed that full-length Hrq1 can bind a poly(dT) ssDNA of 30 nt in length (Fig. 2B). In contrast, however, Hrq1ΔN displayed only modest (<5%) ssDNA binding at the highest concentration of protein tested (5 nM). As with the ATPase assays above, these data again suggest an important role for the Hrq1 N-terminus in ssDNA binding. Indeed, the isolated NTD displayed more ssDNA binding than Hrq1ΔN (Fig. 1C).

By definition, the core motor domains of DNA helicases interact with DNA during unwinding, but the data presented here indicate that Hrq1 has a secondary ssDNA binding site in its NTD that increases the affinity of the protein for its nucleic acid substrate (Fig. 2B and 2C) and coordinates ssDNA binding with ATP hydrolysis (Fig. 2A). This is similar to the NTD of RECQL4, which is responsible for DNA interactions that drive ssDNA strand annealing and DNA strand exchange (Macris, Krejci et al. 2006, Xu and Liu 2009, Keller, Kiosze et al. 2014, Rogers and Bochman 2017, Rogers, Sausen et al. 2019). Although Hrq1 does not have strand annealing or exchange activity (Rogers and Bochman 2017), it may still utilize its NTD – and perhaps the natively disordered portions thereof – to bind to its wide variety of high-affinity substrates like G-quadruplex (G4) structures, D-loops, and RNA-DNA hybrids (Rogers, Wang et al. 2017, Nickens, Rogers et al. 2018). This would again parallel the DNA structure binding displayed by the RECQL4 NTD (Keller, Kiosze et al. 2014, Marino, Mojumdar et al. 2016, Papageorgiou, Pospisilova et al. 2023).

### *In vivo* roles of the NTD

Fibroblasts from Rothmund-Thomson syndrome patients with *recql4* mutations display sensitivity to drugs that induce DNA inter-strand crosslinks (ICLs) but not other types of DNA damage (Jin, Liu et al. 2008), suggesting a role for the helicase in ICL repair. This function appears to be conserved in Hrq1 because we have shown that it functions in the Pso2 ICL repair pathway in yeast (Bochman, Paeschke et al. 2014, Rogers, van Kessel et al. 2014, Rogers, Wang et al. 2017, Rogers, Lee et al. 2020, Rogers, Simmons III et al. 2020). To determine if the Hrq1 NTD is needed for its ICL repair activity, we engineered the *hrq1ΔN* allele in *S. cerevisiae* and exposed the cells to the ICL-inducing agent DEB.

As shown in Figure 3, *hrq1Δ* cells are sensitive to DEB, and the catalytically inactive allele of *HRQ1* (*hrq1-K318A*) displays even greater sensitivity. We have observed this phenomenon before (Bochman, Paeschke et al. 2014, Rogers, Wang et al. 2017, Rogers, Lee et al. 2020) and note that Hrq1-K318A retains DNA binding activity *in vitro* (Nickens, Rogers et al. 2018) and is produced at wild-type levels *in vivo* (Bochman, Paeschke et al. 2014). Thus, we hypothesize that the mutant protein acts as a dominant negative by being recruited to sites of ICL damage and then blocking compensatory pathways from accessing the lesions for repair. In contrast to *hrq1-K318A*, the *hrq1ΔN* mutant phenocopies the null (Fig. 3). This corresponds with the data presented above, showing that recombinant Hrq1ΔN is defective for DNA binding (Fig. 2B and 2C). We hypothesize that the Hrq1ΔN protein does not interact with ICL substrates *in vivo*, explaining why this allele is as ineffective in ICL repair as the *hrq1Δ* strain.

### The genetic interactome of *hrq1ΔN*

To gain further insight into the role(s) of the Hrq1 NTD *in vivo*, we performed SGA analysis of the *hrq1ΔN* allele by crossing it to the *S. cerevisiae* single-gene deletion (Giaever and Nislow 2014) and TS mutant collections (Kofoed, Milbury et al. 2015). Because we previously performed identical analyses for *hrq1Δ* and *hrq1-K318A* strains (Rogers, Sanders et al. 2020, Sanders, Nguyen et al. 2020), we can compare the data for differential interactions that may explain the distinct phenotype of our *hrq1* mutants. Here, we found 286 synthetic genetic interactions with *hrq1ΔN* that were nearly evenly split between the deletion (151) and TS collections (135). The split was also similar between negative (81 in the deletion collection, 76 in the TS collection) and positive genetic interactions (70 in the deletion collection, 59 in the TS collection), and both types of interactors are listed in Table 1.

**Table 1.** Synthetic genetic interactions with *hrq1ΔN*.

| Negative interactions<sup>a</sup> | Positive interactions |
| --- | --- |
| Gene deletions |  |
| YSC83, SLX9, GIC1, MRM2, SPO16, UTH1, TOM70, SWC5, PHO5, YME1, YJU3, MDM35, RPL6B, YUR1, STE50, MUP3, HAP2, HAP3, PCK1, BEM1, DAK1, TIM18, ASK10, UBP1, MBR1, TCO89, RAD4, KCH1, SWI4, AIM4, YJL213W, ENT3, MIR1, PET127, CDC73, GPR1, ELM1, RAD14, IRA2, BDF2, PHB2, MLS1, PMA2, VAM6, CSE2, MNN11, BTS1, IMG2, YPR084W, YML090W, TGS1, MDH3, GRR1, HTD2, SPO20, COX10, YJL175W, IXR1, PEF1, YHK8, QCR9, KGD2, YOS9, KAP114, PET494, UPS1, YMR245W, OAR1, LTE1, RPL43B, PAC10, YBR238C, CTI6, LPD1, FPR2, LIP5, LDB19, YJR120W, SAP190, SIN3, COX23 | INP2, RPS27B, OLA1, DGA1, YML012C-A, YLR338W, AAH1, SWI6, BNA6, DIA2, ATG22, TSR2, ERG3, UBS1, VRP1, ILM1, ADE2, ACB1, PEX12, TSR3, VPS9, ARF1, RPO41, NHP10, YHM2, SIM1, MSH4, SHE4, OST4, VPS4, TRM44, SPR6, PDE1, RPN4, ART5, VID30, YGL024W, EDC1, CSM1, ALT1, YBL094C, SAC1, YHR180W, NDE1, ARC1, YAR1, YDL062W, FEN1, YBR016W, YMR031W-A, GRX2, MSB4, YMR294W-A, IES5, YJL206C-A, EOS1, VAM7, PKP2, HXT2, KAR5, CAT2, MFT1, MKS1, RTG1, ICE2, ARA2, RPS9B, RTG3, RTG2, YGL214W, |
| Temperature-sensitive alleles |  |
| yef3-f650s, mob2-8, efb1-4, mob2-38, taf2-1, mob2-36, mob2-14, mob2-28, cse2, mob2-34, med7-163, cdc31-5, mob2-40, mob2-20, mob2-11, mps1-1, mob2-19, smc2-8, mob2-22, pol1-17, yhr172w-ph, ycs4-1, smd1-1, rlp7-ph, yhc1-2, tub2-443, dbp5-1, rox3-182, pse1-34, sec63-1, mcm1-ph, cdc21-1, mas2-10, arp3-g15c, mob2-24, eaf7, puf6, spt3, htd2, apq12, bbp1-1, cdc20-1, nup57-e17, lst8-6, yil061c-ph, neo1-2, cof1-5, pob3-q308k, ndc1-4, ykr022c-ph, lip5, ipi1-ph, rrp4-1, yil106w-ph, sgf73, dfr1-td, fip1-ph, srb6-ph, lst8-15, tfg2-ph, slm3, nab3-11, pam16-3, pre7-ph, lge1, cct4-1, swd2-ts1, spc29-3, stb5, nut2-ts, dop1-1, arp3-f306g, pth1, cmd1-3, rps22b, qcr2 | gpi12-ph, myo2-14, sts1-ph, ymr134w-ph, slx9, sec24-2, sec27-1, sec23-1, gtr1, pds5-1, ies5, act1-129, vam6, ykt6-ph, srv2-2, orc2-4, sed5-1, act1-112, rpf2-ph, stt3-2, stu2-10, pfy1-4, rvb2-ph, bet3-ph, orc1-ph, sur1, act1-2, yhc1-6, ies1, stt3-1, gpi10-ph, gaa1-ts, rpc40-v78r, pbn1-ph, smc4-1, nhp10, sda1-2, brn1-9, spt5-194, rsp5-sm1, act1-155, nse4-ph, act1-3, ebp2-1, cdc1-4, dpb11-1, cdc11-5, arp3-d11a, rrn7-ph, sfi1-7, fal1-1, spt6-14, cdc2-7, cdc48-1, ydl003w-ph, apc11-22, arp3-g302y, las17-1, mps3-1 |

<sup>a</sup> Negative genetic interactors are listed from largest absolute value of the SGA score to the smallest, but positive interactors are listed from the smallest to the largest.

Figure 4 shows the frequency distribution of the SGA scores for the screens of both collections as violin plots (Fig. 4A), with separate box blots for the negative (Fig. 4B) and positive (Fig. 4C) synthetic genetic interactions, where the outliers are denoted with individual points. These outliers are the mutants with the statistically strongest phenotypes. As depicted in Figure 4 though, the vast majority of the synthetic phenotypes were mild increases or decreases in growth of the double-mutants compared to the parental strains. This is similar to the SGA analyses of *hrq1Δ* and *hrq1-K318A*, as well as their equivalent alleles of the Sgs1 helicase (Sanders, Nguyen et al. 2020). Screening double mutants of *hrq1* and *sgs1* by SGA lead to a synergistic increase in the number of triple mutant strains with strong growth phenotypes, so it would be interesting to also, for instance, cross *hrq1ΔN sgs1Δ* and *hrq1ΔN sgs1-K706A* strains to the deletion collection for analysis.

**Figure 4.**
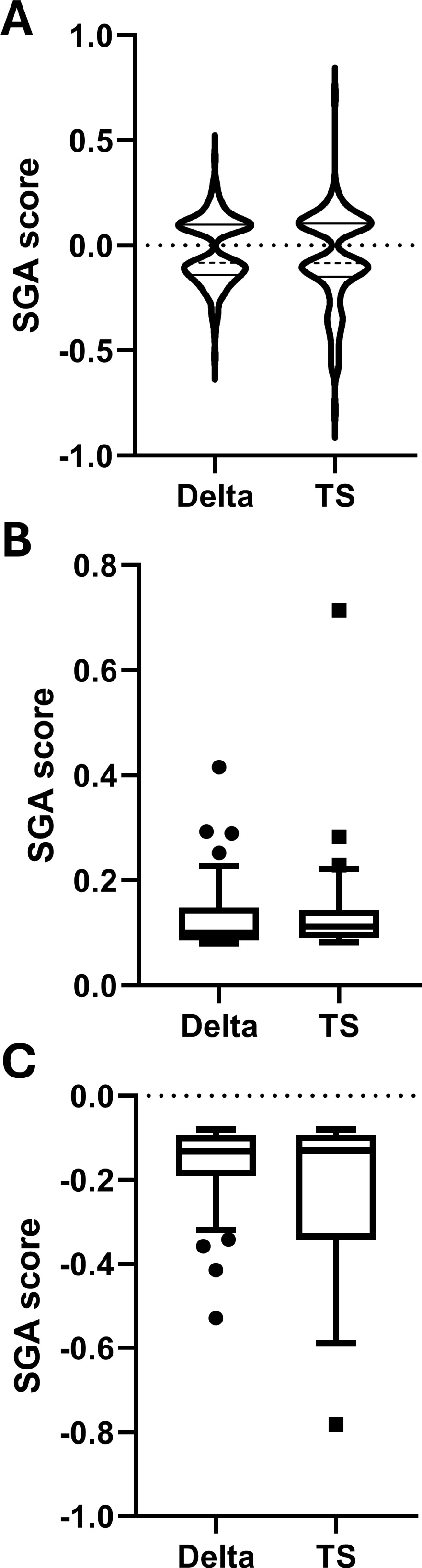
Analysis of the distribution and magnitudes of the synthetic genetic interactions with *hrq1ΔN*. **A)** Violin plots of the synthetic genetic interactions with the single-gene deletion collection (Delta) and TS collection. The median values are denoted with dashed lines, and the quartiles are shown as solid lines. The SGA data are also shown in separate box and whisker plots drawn using the Tukey method for the positive **(B)** and negative **(C)** interactions with the deletion and TS collections. The individually plotted points outside of the inner fences represent outliers (*i.e.*, interactions with mutants yielding the strongest SGA scores) and correspond to alleles whose SGA score is greater than the value of the 75^th^ quartile plus 1.5 times the inter-quartile distance (IQR) for positive interactions and alleles whose SGA score is less than the value of the 25^th^ quartile minus 1.5IQR for negative interactions. The significant differences between SGA data sets discussed in the main text were calculated using the Kruskal-Wallis test and Dunn’s multiple comparisons test.

### *hrq1ΔN* interactions

Regardless, combining the deletion and TS data, loss of the Hrq1 NTD yielded strong positive genetic interactions with seven genes: *RPS9B*, *RTG3*, *RTG2*, *YGL214W*, *ARP3*, *LAS17*, and *MPS3*. The *RPS9B* gene encodes a protein found in the small ribosomal subunit (Mager, Planta et al. 1997), and Hrq1 has previously been linked to ribosome assembly and maturation at the genetic and physical interactome levels (Rogers, Sanders et al. 2020). Rtg2 regulates the subcellular localization of Rtg3 and communicates mitochondrial dysfunction to the nucleus (Liao and Butow 1993, Sekito, Thornton et al. 2000). Hrq1 and RECQL4 are the only known RecQ family helicases with both nuclear and mitochondrial roles, though their functions in mitochondria remain unclear (Croteau, Rossi et al. 2012, Koh, Chong et al. 2015). It is tempting to speculate that Hrq1 is also involved in this nuclear-mitochondrial communication pathway or acts downstream of the communication to help maintain the stability of the nuclear and/or mitochondrial genomes. Arp3 and Las17 are interacting proteins that regulate actin filaments (Machesky and Gould 1999, Madania, Dumoulin et al. 1999). Although it is unclear what Hrq1 may have to do with the cytoskeleton, *arp3* mutation also decreases telomere length (Ungar, Yosef et al. 2009), and Hrq1 has multiple links to telomere biology (Bochman, Paeschke et al. 2014, Rogers, Wang et al. 2017, Nickens, Rogers et al. 2018, Nickens, Sausen et al. 2019, Nickens and Bochman 2021, Nickens, Feng et al. 2024). Therefore, synthetic genetic interaction between *hrq1ΔN* and *arp3Δ* may be related to their roles in telomere maintenance. However, human RECQL4 is a microtubule-interacting protein (Yokoyama, Moreno-Andres et al. 2019), so an interaction between Hrq1 and the cytoskeleton cannot be ruled out. In support of this is the fact that Mps3 localizes to telomeres during meiosis, and deletion of *MSP3* likewise synergizes with *hrq1ΔN.* Finally, *YGL214W* is a dubious ORF that likely does not encode a functional protein (https://www.yeastgenome.org/locus/S000003182). However, the *ygl214wΔ* allele in the deletion collection also disrupts the nearby *SKI8* gene (Kushner, Lindenbach et al. 2003), and Ski8 is required for DSB repair during meiosis (Arora, Kee et al. 2004).

Similar to the above, the combined SGA data revealed five strong negative genetic interactions (*YSC83*, *SLX9*, *GIC1*, *MRM2*, and *YEF3*), and common themes emerge. For instance, Slx9 is involved in ribosome biogenesis (Bax, Raue et al. 2006), and Yef3 is a translation elongation factor (Chakraburtty and Triana-Alonso 1998), so perhaps their double mutants with *hrq1ΔN* are related to Hrq1’s putative role in ribosome biology (Rogers, Sanders et al. 2020). That said, Slx9 is also a G4 structure-binding protein (Gotz, Pandey et al. 2019), and G4 DNA is a preferred substrate for Hrq1 *in vitro* (Rogers, Wang et al. 2017). Therefore, the negative genetic interaction between *hrq1ΔN* and *slx9Δ* could be due to their roles in DNA secondary structure maintenance. In contrast, Ysc83 and Mrm2 are mitochondrial proteins (Pintard, Bujnicki et al. 2002, Sickmann, Reinders et al. 2003), and the negative genetic interactions here again point toward Hrq1’s undefined role(s) in mitochondria. *GIC1* encodes a protein that regulates mitotic exit, with links to regulation of the actin cytoskeleton (Hofken and Schiebel 2004), but as stated above, how actin is connected to Hrq1 and its NTD remains unknown.

### Cellular pathways affected by *hrq1ΔN*

Next, we used Gene Ontology (GO) Term enrichment analysis to determine which pathways in yeast are affected by *hrq1ΔN* and mutation in other genes. Table 2 shows the top 10 GO Terms and their associated genes related to negative genetic interactions with *hrq1ΔN*. The GO Terms include transcription by RNA polymerase II, regulation of organelle organization, DNA repair, chromatin organization, mitotic cell cycle, peptidyl-amino acid modification, cytoskeleton organization, mitochondrion organization, organelle fission, and response to chemical. Similarly, Table 3 collates the top 10 GO Terms related to positive genetic interactions with *hrq1ΔN* (mitotic cell cycle, cytoskeleton organization, chromatin organization, lipid metabolic process, organelle fission, DNA repair, rRNA processing, transcription by RNA polymerase II, response to chemical, and regulation of organelle organization), which overlap by 80% with the GO Terms for negative interactions.

**Table 2.** Gene Ontology (GO) Term enrichment of negative genetic interactors with *hrq1ΔN*.

| GO Term (GO ID) | Genes annotated to the GO Term<sup>a</sup> | GO Term usage in gene list | Genome frequency of use |
| --- | --- | --- | --- |
| transcription by RNA polymerase II<br>(GO:0006366) | CDC73, CSE2, EAF7, IXR1, MCM1, MED7, NAB3, NUT2, POB3, RAD4, ROX3, SGF73, SIN3, SPT3, SRB6, STB5, SWI4, TAF2, TFG2, YBL021C, YGL237C | 21 of 138 genes,<br>15.22% | 490 of 6488 annotated genes,<br>7.55% |
| mitotic cell cycle<br>(GO:0000278) | CDC20, ELM1, GIC1, GRR1, LTE1, MCM1, MOB1, MPS1, PEF1, PSE1, SAP190, SIN3, SMC2, SPC29, SWI4, TUB2, YCS4 | 17 of 138 genes,<br>12.32% | 330 of 6488 annotated genes,<br>5.09% |
| cytoskeleton organization<br>(GO:0007010) | ARP3, BBP1, CDC31, CMD1, COF1, EFB1, ELM1, ENT3, GIC1, LST8, MPS1, NDC1, SPC29, SPC97, TUB2 | 15 of 138 genes,<br>10.87% | 252 of 6488 annotated genes,<br>3.88% |
| response to chemical<br>(GO:0042221) | ASK10, CMD1, GPR1, GRR1, IXR1, LDB19, STB5, STE50, TIM18, YHK8, YOS9 | 11 of 138 genes,<br>7.97% | 408 of 6488 annotated genes,<br>6.29% |
| transmembrane transport (GO:0055085) | ASK10, KCH1, MIR1, MUP3, PAM16, PMA2, SEC63, TOM70, YHK8, YME1 | 10 of 138 genes,<br>7.25% | 278 of 6488 annotated genes,<br>4.28% |
| regulation of cell cycle<br>(GO:0051726) | CDC20, GIC1, GRR1, LTE1, MOB1, MPS1, PSE1, SMC2, SPO16, YCS4 | 10 of 138 genes,<br>7.25% | 245 of 6488 annotated genes,<br>3.78% |
| organelle fission<br>(GO:0048285) | CDC20, GIC1, LTE1, MOB1, MPS1, PSE1, SMC2, SPO16, TUB2, YCS4 | 10 of 138 genes,<br>7.25% | 225 of 6488 annotated genes,<br>3.47% |
| meiotic cell cycle<br>(GO:0051321) | CDC20, LGE1, MPS1, POL1, SIN3, SMC2, SPO16, SPO20, TGS1, YCS4 | 10 of 138 genes,<br>7.25% | 289 of 6488 annotated genes,<br>4.45% |
| cellular respiration<br>(GO:0045333) | KGD2, LPD1, MBR1, OAR1, QCR2, QCR9, YBL021C, YBR238C, YGL237C, YJR120W | 10 of 138 genes,<br>7.25% | 86 of 6488 annotated genes,<br>1.33% |
| nucleus organization<br>(GO:0006997) | APQ12, CMD1, CTI6, NDC1, NUP57, SIN3, SMC2, TGS1, YCS4 | 9 of 138 genes,<br>6.52% | 99 of 6488 annotated genes,<br>1.53% |

<sup>a</sup> Three negative genetic interactors (*YML090W*, *YJL175W*, and *YMR245W*) represent dubious ORFs, and 11 (*FPR2*, *YPR084W*, *YSC83*, *DFR1*, *IRA2*, *AIM4*, *YJL213W*, *KAP114*, *PAC10*, *COX10*, and *PET127*) could not be mapped to terms in the current GO set or are annotated to the root node for the set being used. Thus, these 14 genes do not appear in the GO analysis.

**Table 3.** Gene Ontology (GO) Term enrichment of positive genetic interactors with *hrq1ΔN*.

| GO Term (GO ID) | Genes annotated to the GO Term<sup>a</sup> | GO Term usage in gene list | Genome frequency of use |
| --- | --- | --- | --- |
| mitotic cell cycle<br>(GO:0000278) | ACT1, BRN1, CDC11, CDC48, DIA2, DPB11, MCD1, MPS3, MYO2, PDS5, PFY1, SDA1, SFI1, SMC4, SPT6, STU2, SWI6, VRP1 | 18 of 113 genes,<br>15.93% | 330 of 6488 annotated genes,<br>5.09% |
| cytoskeleton organization<br>(GO:0007010) | ACT1, ARP3, CDC48, ICE2, LAS17, MPS3, MSB4, MYO2, PFY1, RSP5, SFI1, SHE4, SRV2, STU2, VRP1 | 15 of 113 genes,<br>13.27% | 252 of 6488 annotated genes,<br>3.88% |
| chromatin organization<br>(GO:0006325) | CSM1, DIA2, GTR1, IES1, MPS3, NHP10, ORC1, ORC2, PDS5, RSP5, RVB2, SMC4, SPT5, SPT6 | 14 of 113 genes,<br>12.39% | 256 of 6488 annotated genes,<br>3.95% |
| lipid metabolic process<br>(GO:0006629) | ACB1, ARF1, CDC1, DGA1, ERG3, GPI10, GPI12, PBN1, RSP5, SAC1, SUR1, VPS4, YMR134W | 13 of 113 genes,<br>11.50% | 296 of 6488 annotated genes,<br>4.56% |
| organelle fission<br>(GO:0048285) | ARF1, BRN1, CSM1, EBP2, MCD1, MPS3, MSH4, PDS5, SMC4, SRV2, SWI6 | 11 of 113 genes,<br>9.73% | 225 of 6488 annotated genes,<br>3.47% |
| DNA repair<br>(GO:0006281) | ACT1, CDC1, DPB11, MCD1, NHP10, NSE4, PDS5, POL3, RAD27, RPN4, SPT5 | 11 of 113 genes,<br>9.73% | 269 of 6488 annotated genes,<br>4.15% |
| rRNA processing<br>(GO:0006364) | EBP2, FAL1, RPF2, RPS27B, RPS9B, RSP5, RVB2, SLX9, SPT5, TSR2, TSR3 | 11 of 113 genes,<br>9.73% | 327 of 6488 annotated genes,<br>5.04% |
| transcription by RNA polymerase II<br>(GO:0006366) | MKS1, RPN4, RSP5, RTG1, RTG2, RTG3, RVB2, SPT5, SPT6, SWI6, YML062C | 11 of 113 genes,<br>9.73% | 490 of 6488 annotated genes,<br>7.55% |
| response to chemical<br>(GO:0042221) | CDC48, EOS1, GRX2, GTR1, PBN1, RPN4, RSP5, RTG1, RTG3, YAR1 | 10 of 113 genes,<br>8.85% | 408 of 6488 annotated genes,<br>6.29% |
| regulation of organelle organization<br>(GO:0033043) | ARF1, CDC48, LAS17, PFY1, RSP5, SEC23, SRV2, SWI6, VRP1 | 9 of 113 genes,<br>7.96% | 227 of 6488 annotated genes,<br>3.50% |

<sup>a</sup> Eight positive genetic interactors (*YML012C-A*, *YLR338W*, *YGL024W*, *YBL094C*, *YDL062W*, *YMR031W-A*, *YMR294W-A*, and *YGL214W*) represent dubious ORFs, and seven (*CAT2*, *YBR016W*, *ARA2*, *YHR180W*, *PDE1*, *NCE101*, and *PKP2*) could not be mapped to terms in the current GO set or are annotated to the root node for the set being used. Thus, these 15 genes do not appear in the GO analysis.

These results make sense with regard to, for instance, Hrq1’s connections to transcription (Rogers, Sanders et al. 2020), DNA ICL repair (Bochman, Paeschke et al. 2014, Rogers, Wang et al. 2017, Rogers, Lee et al. 2020), and mitochondrial localization (Koh, Chong et al. 2015). Similarly, they reinforce the strong synthetic interactions described above (*e.g.*, cytoskeleton organization and the synergism between *hrq1ΔN* and *arp3*, *las17*, and *gic1*; Fig. 4). Therefore, investigating the genetic interactions between *hrq1ΔN* and other *hrq1* alleles should be fertile ground for future research into multiple physiological functions for this RecQ family helicase.

### How do the synthetic genetic interactions with *hrq1ΔN* compare to those with *hrq1Δ* and *hrq1-K318A*?

Our *in vitro* biochemistry demonstrates that Hrq1ΔN is still a catalytically active protein (Fig. 2), suggesting that it could perform some of the functions of Hrq1 *in vivo.* However, cells expressing *hrq1ΔN* display *hrq1Δ*-levels of DNA ICL sensitivity (Fig. 3). This begs the question: how do the synthetic genetic interactions with *hrq1ΔN* uncovered here compare to those published for *hrq1Δ* and *hrq1-K318A*? Analyzing all combinations of positive and negative synthetic genetic interactions shows that there is similar overlap between mutants from the deletion and TS collections with all of the *hrq1* alleles investigated (Fig. 5). In general, the genetic interactions overlapped more with TS alleles, with 53-77% of genetic interactions shared between *hrq1Δ* or *hrq1-K318A* and the *hrq1ΔN* data set. The interactions with the deletion collection were more varied, with 33-58% overlap comparing *hrq1ΔN* to *hrq1Δ* and *hrq1-K318A*. These data are similar to those for comparisons between *hrq1Δ* and *hrq1-K318A* themselves (Sanders, Nguyen et al. 2020).

**Figure 5.**
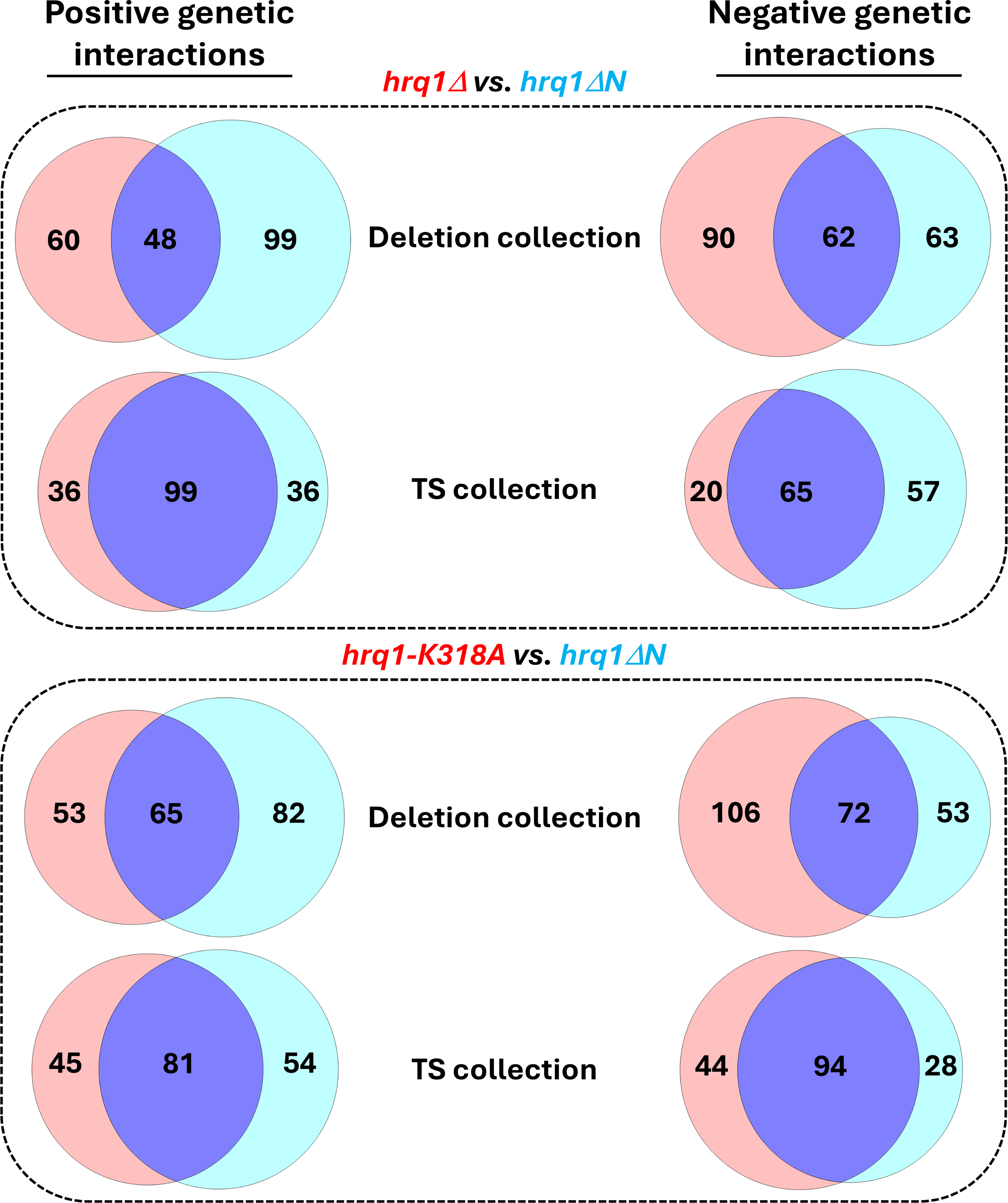
Venn diagrams of the shared synthetic genetic interactions displayed by *hrq1Δ* and *hrq1ΔN* (top) and *hrq1-K318A* and *hrq1ΔN* (bottom). The numbers of unique and overlapping interactions are labeled. The circles and their percent overlap were drawn to scale using Venn Diagram Plotter software (v. 1.6.7458.27187).

## CONCLUSIONS AND PERSPECTIVES

Here, we have reported a comprehensive set of synthetic genetic interactions between nearly all of the genes in the *S. cerevisiae* genome and the NTD-truncated *hrq1ΔN* allele of the Hrq1 helicase, a homolog of disease-linked human RECQL4. The data sets included herein supplement the existing sets of known *hrq1Δ* and *hrq1-K318A* interactions (Rogers, Sanders et al. 2020, Sanders, Nguyen et al. 2020), expanding the genetic interactome landscape of *hrq1*. As with the deletion and catalytically inactive mutants, it is clear that *hrq1ΔN* genetically interacts with genes in multiple important pathways in yeast that underpin the maintenance of genome integrity. As such, these SGA analyses have also helped us to generate testable hypotheses that are driving our ongoing and planned research. As with *hrq1Δ* and *hrq1-K318A*, the *hrq1ΔN* genetic interactome again suggests links to transcription, suggesting that RECQL4 may also functionally interact with the transcription machinery as previously established for human RECQL5 and RNA polymerase II (Aygun, Svejstrup et al. 2008, Izumikawa, Yanagida et al. 2008, Saponaro, Kantidakis et al. 2014). Indeed, we are currently investigating potential roles for Hrq1 in transcription and are establishing tools to extend this work to the *Drosophila melanogaster* RECQL4 homolog (Capp, Wu et al. 2009). It is our hope that the data provided here will spur additional work in the field on RecQ4 subfamily helicases to finally elucidate the roles of these enzymes in genome integrity.

## DATA AVAILABILITY STATEMENT

The data underlying this article are available in the article, in its online supplementary material, or will be shared upon reasonable request to the corresponding author.

## ACKNOWLEDGMENTS

We thank Amy Caudy for sharing plasmids, the University of Toronto for performing the SGA analyses, Michael Costanzo and members of the Boone lab for help with data collection and interpretation, and members of the Bochman lab for critically reading this manuscript.

